# Nuclear Ribosomal Internal Transcribed Spacer 1 (ITS1) variation in Lady Bird Beetles (Coccinellidae)

**DOI:** 10.1101/2022.05.11.491562

**Authors:** A. G. B. Aruggoda, Shun-Xiang Ren

## Abstract

The seventeen taxa of the family Coccinellidae were analyzed by sequencing the Internal Transcribed Spacer 1 region followed by analyzing the presence of significantly simple motifs (SSMs), to investigate the presence of functionally important areas in the gene if any and to find phylogenetic inference of the ITSI region between sequenced species. Length comparisons among the ITS 1 region indicated the absence of intraspecific variability. Further, length differences observed were among *Adalia bipunctata* showing a 3 bp maximum size difference and *Psyllobora vigintiduopunctata* with 1 bp size difference. The base composition showed highly similar values of A, T, G and C. Except *Harmonia axyridis, Phyllobora vigintiduopunctata, Chilocorus renipustulatu, Lemnia duvauceli* all other species showed significant number of simple repetitions. *Adalia bipunctata* showed the highest simple repetition with the RSF value of 1.4440. Thirteen ladybird beetles had higher SSM regions. *Adalia bipunctata, Adalia decempunctata, Exochomus quadripustulatus* shows high accumulation of AT repeats in 5 prime end flanking region and GC repeats in 3 prime end flanking region. *Amida nigropectoralis, Amida quingquefasiata, Menochilus sexmaculata, Phaenochilus mesternalis, Coccidula rufa, Platynaspis luterubra s*hows a higher concentration of GC repeats towards the 3 prime end flanking regions. Phylogenetic analysis revealed that the tribe Psylloborini of the subfamily Coccinellidae was monophyletic. The members of the Subfamily Chilocorinae first associate with the tribes of subfamily Coccinellidae and then relate with the tribe Scymnini. *Exochomus quadripustulatus* which shows the highest length variably showed a relationship with *Phaenochilus mesternalis* 77 bootstrap and 84 posterior probabilities.

## 1.0 Introduction

The first and second internal transcribed spacers (ITS1 and ITS2) are located between 18S and 28S gene regions whereas ITS 1 is located between 18S and 5.8S and ITS 2 is located between 5.8S and 28S rRNA gene coding regions. Eukaryotes’ ribosomal RNA genes (rDNA) are organized in clusters of tandem repeats, each consisting of coding regions with transcribed and non-transcribed spacers. The gene region starts with the external transcribed spacer (ETS) at the 5’ end, downstream by the 18S rRNA gene, to the internal transcribed spacer 1 (ITS1), and the 5.8S rRNA gene region followed by, the internal transcribed spacer 2 (ITS2) and the 28S rRNA gene, terminating with the intergenic spacer (IGS) (Gerbi,1985). The 5.8S gene exhibits a slow rate of evolutionary change, however, the level of sequence variation of the spacers is higher (Hillis and Dixon, 1991) therefore possible to infer phylogenetic relationships at the higher taxonomic levels such as subfamily, and family levels (González et al., 1990; Vogler and De Salle, 1994; Coleman and Vacquier, 2002). Generally, several copies are found per genome, at one or more chromosomal sites termed nucleolus organizer regions (NORs) (Long and Dawid, 1980; Mindell and Honeycutt, 1990) which are composed of 18S, 5.8S and 28S ribosomal RNA; separated by two intergenic spacers and an external transcribed spacer. These gene regions have become popular marker among researchers to study phylogenetic relationships among closely related species of animals, plants, and fungi specially because it exhibits a high rate of evolution due to their noncoding structure and can be easily multiplied via PCR from almost any taxon, using conserved primers (Hillis and Dixon, 1991; Bakker, et al., 1995; Buckler and Holtsford., 1996; Gouliamova and Hennebert 1998; Gouliamova et al, 1998; von der Schulenburg et al., 2001). Further, intraindividual variation is shown by the ITS region therefore several researchers have been interpreted accurately the information variation among repeats within genomes in a range of taxa (Vogler and De Salle, 1994; Tang et al., 1996; Harris and Crandall, 2000; Gandolfi et al., 2001; Hartmann et al., 2001; Mayol and Rosselló, 2001; von der Schulenburg et al., 2001; Leo and Barker, 2002). In addition, ITS1 evolution has shown internal repetition, leading to size variation. The repetition includes comparatively long repeat units, e.g., in trematodes (Platyhelminthes) and dipterans (Paskewitz, et al., 1993; O’Kane et al. 1996; Tang et al. 1996; van Herwerden, et al., 1998, 1999), or commonly, ‘‘simple’’ repetitive sequence motifs, of various arthropods and in humans (Gonzalez et al. 1990; Kwon and Ishikawa 1992; Wesson, et al., 1992; Wesson et al. 1993; Kuperus and Chapco 1994; Vogler and De Salle 1994; McLain et al. 1995; Miller et al., 1996; Fenton, et al., 1997; Remigio and Blair 1997; Kumar, et al., 1999; Harris and Crandall 2000).

The continuous molecular processes such as replication slippage, unequal crossing over, and biased gene conversion, which are involved in the generation of length variation and/or lead to concerted evolution are common in the repetitive sequences (Dover 1982; Levinson and Gutman 1987; Elder and Turner 1995). ITS1 evolution may further be affected by the extent of homogenization between repeat units of the ribosomal cistron. Inefficient homogenization between repeats may result in intraindividual ITS1 heteroplasmy, as hypothesized for some insects, crustaceans, and trematodes (Wesson, et al., 1992; Vogler and De Salle 1994; van Herwerden, et al., 1998; Harris and Crandall 2000). Consequently, the evolution of these spacers seems to be characterized by various factors that are poorly understood. This information definitely should have a value in understanding the evolution of Internal transcribed spacer region giving a value to the ITS as a reliable marker in phylogenetic studies. In 2001, von der Schulenburg et al., discovered the extreme length and length variation of ITS1 in ladybird beetles (Coleoptera: Coccinellidae) giving the researchers an ideal opportunity to expand the current knowledge on the evolutionary dynamics of these spacers. Thereafter, no one gave attention to understand the evolutionary relationships among extreme length, simple repeats and the evolutionary patterns among the Coccinellidae. Expanding this understanding is worth enough to the current world scenario of managing environmentally friendly pests, by that there is a tendency to develop predatory incidences of the agriculturally important Coccinellidae species. The present study aimed to study three major areas of internal transcribed spacer 1 region of Coccinellidae; first to investigate the presence of significantly simple motifs (SSMs) other than long repetitive units; second to investigate the presence of functionally important areas in the gene if any; third to find the phylogenetic inference of the ITSI region between sequenced species.

## 2.0 Methodology

### 2.1 Taxon sampling

The seventeen taxa represented five proposed subfamilies and seven proposed tribes were analyzed by sequencing Internal Transcribed Spacer 1 region (Table 01). The taxon sampling was carried out as per the classification outlined by Pang et al., 1979 and Kovář in 1996. The samples of species belonging to the subfamilies of Coccinellidae: Coccinellinae, Scymninae, Coccidulinae, Chilochorinae and Ortaliinae (present taxonomic position from Kovář 1996) and tribes, Coccinellini, Psylloborini, Ortaliini, Coccidulini, Scymnini, Platynaspini and Chilocorini.

**Table 01.** Taxa studied and their Genbank accession numbers.

| Family | Subfamily | Tribe | Species | Collection locality | Length of ITS I region (bp) | Total length (bp) | Genbank Accession No. |
| --- | --- | --- | --- | --- | --- | --- | --- |
| Coccinellidae | Coccinellinae | Psyllobroini | *Halysia sanscrita* | Ailaoshan, Yunnan | 951 | 1107 | ON318315 |
| Coccinellidae |  | Psyllobroini | *Psyllobora vigintiduopunctata* | Taibai, Shaanxi | 992 | 1148 | ON318314 |
|  |  |  |  | Berlin, German | 992 | 1148 | AJ272149 |
| Coccinellidae |  | Coccinellini | *Adalia bipunctata* | Zhongdian, Yunnan | 1515 | 1671 | ON318312 |
|  |  |  |  | Brelin, German | 1515 | 1671 | AJ272141 |
|  |  |  |  | Cambridge,UK | 1512 | 1668 | AJ272140 |
|  |  |  |  | Moscow, Russia | 1514 | 1670 | AJ272139 |
| Coccinellidae |  | Coccinellini | *Adalia decempunctata* | Ailaoshan, Yunnan | 1401 | 1557 | ON318313 |
|  |  |  |  | Brelin, German | 1401 | 1557 | AJ272148 |
| Coccinellidae |  | Coccinellini | *Coccinella septempunctata* | Ailaoshan, Yunnan | 911 | 1067 | ON318311 |
|  |  |  |  | Berlin, German | 911 | 1067 | AJ272143 |
| Coccinellidae |  | Coccinellini | *Harmonia axyridis* | Xishan, Yunnan | 1188 | 1344 | ON318310 |
|  |  |  |  | Berlin, German | 1188 | 1344 | AJ272146 |
| Coccinellidae |  | Coccinellini | *Micraspis discolor* | Nanling, Gunagdong | 1113 | 1269 | ON318309 |
| Coccinellidae |  | Coccinellini | *Lemnia duvauceli* | Jinghong, Yunnan | 1320 | 1476 | ON318307 |
| Coccinellidae |  | Coccinellini | *Menochilus sexmaculata* | Nanling, Gunagdong | 1008 | 1164 | ON318308 |
| Coccinellidae | Chilocorinae | Chilocorini | *Phaenochilus mesternalis* | Wuzhishan, Hainan | 826 | 982 | ON318317 |
| Coccinellidae |  | Chilocorini | *Chilocorus renipustulatus* | Jinggongshan, Jiangxi | 1100 | 1256 | ON318318 |
|  |  |  |  | Brelin, German | 1100 | 1256 | AJ272147 |
| Coccinellidae |  | Chilocorini | *Exochomus quadripustulatus* | Limushan, Hainan | 2572 | 2728 | ON318316 |
|  |  |  |  | Brelín, German | 2572 | 2728 | AJ272150 |
| Coccinellidae |  | Platynaspini | *Platynaspis luterubra* | Shimentai, Gunagdong | 1042 | 1198 | ON318321 |
|  |  |  |  | Brelín, German | 1042 | 1198 | AJ272144 |
| Coccinellidae | Coccidulinae | Coccidulini | *Coccidula rufa* | Xishan, Yunnan | 791 | 947 | ON318319 |
|  |  |  |  | Brelín, German | 791 | 947 | AJ272145 |
| Coccinellidae |  | Scymnini | *Scymnus suturalis* | Shimentai, Gunagdong | 1124 | 1280 | ON318320 |
|  |  |  |  | Brelín, German | 1124 | 1280 | AJ272142 |
| Coccinellidae |  | Ortaliini | *Amida pectoralis* | Nangling Gunagzhou | 810 | 966 | ON318322 |
| Coccinellidae |  | Ortaliini | *Amida quinquefasciata* | Yingole, Gunagdong | 823 | 979 | ON318323 |
| Chrysomelidae |  |  | *Diabrotica barberi* | Gunagzhou | 616 | 645 | ON318324 |
| Chrysomelidae |  |  | *Cerotoma trifurcata* | Gunagzhou | 616 | 645 | ON318325 |

Sequences of twelve (12) Coccinellidae species; NCBI accession numbers AJ272139 to AJ272150 (von der Schulenburg *et al.*, 2001) were added to table number one (01) for length comparison. The outgroups were selected from the closely-related Chrysomelidae - infraorder. Generally, one specimen per species was used for the DNA extraction.

### 2.2 DNA extraction, PCR amplification, sequencing, and alignment

DNA was extracted from 95% ethanol-preserved specimens using protocol added with proteinase K. The small-bodied specimens were split between the pro-and mesothorax and utilized about 2 mg a part of the thorax tissue for DNA extraction. For larger beetles, a single leg was removed and split along both the femur and tibia to utilize the tissue sample having the same weight. Before dissecting beetles were soaked overnight in 1.5 ml TE buffer (pH 8.0). 2 mg of the tissue was used for DNA extraction. The polymerase chain reaction was carried out with 50 μl PCR reaction volumes. Each 50 μl reaction mixture contained 0.5 μl Taq polymerase - TAKARA (5 units/μl), 5 μl 10x Taq buffer, 5 μl Mgcl (25 mM), 4 μl dNTP mixture (25 mM), 2 μl each primer (10 μMl), 3 μl of DNA extract and 28.5 μl of Doubled distilled water. The primer sequences and annealing temperatures relevant to each primer are given in table 02. PCR products were sequenced from two directions using ABI-3730 DNA sequencer with ABI Prism Big Dye cycle sequencing kit. The genes which were having higher lengths and which could not be sequenced directly were cloned to the pMD18-T vector system. *E. coli* strain TOP-10 was used as the competent cell for cloning. The cloning procedure followed was the same as in part one.

**Table 02:**
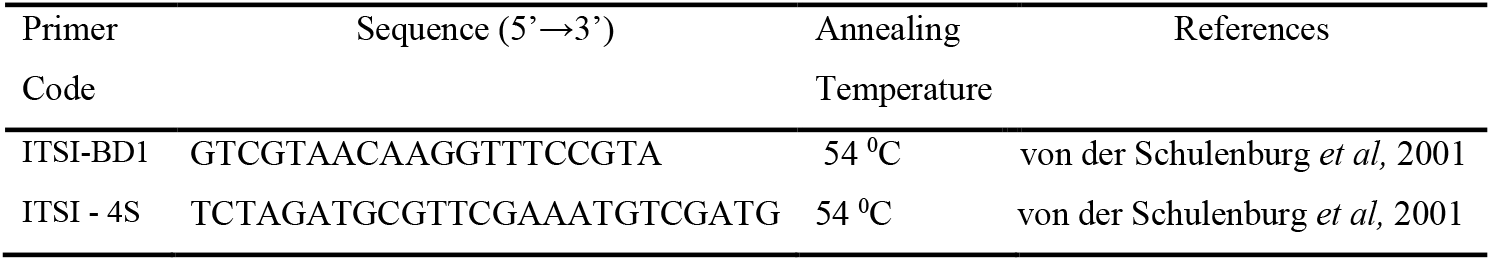
Primers used to amplify ITSI region with partial sequences of 18S and 5.8S.

**Table 03:** Relative simplicity factors (RSF), and presence of significant simple motifs.

| Species | ITS1 spacer length (bp) | RSF | Presence of simple motifs (SSM) |
| --- | --- | --- | --- |
| *Adalia bipunctata* | 1515 | 1.4440 | significant |
| *Adalia decempunctata* | 1401 | 1.2135 | significant |
| *Menochilus sexmaculata* | 1008 | 1.1427 | significant |
| *Coccinella septempunctata* | 911 | 1.2159 | significant |
| *Harmonia axyridis* | 1188 | 1.0833 | Non - significant |
| *Lemnia duvauceli* | 1320 | 1.0940 | Non - significant |
| *Micraspis discolor* | 1112 | 1.1232 | Non - significant |
| *Phyllobora vigintiduopunctata* | 992 | 1.0959 | Non - significant |
| *Halysia sanscrita* | 951 | 1.0891 | significant |
| *Chilocorus renipustulatus* | 1100 | 1.0201 | Non - significant |
| *Phaenochilus mesternalis* | 826 | 1.0937 | Non - significant |
| *Platynaspis luterubra* | 1042 | 1.1245 | significant |
| *Exochomus quadripustulatus* | 2572 | 1.1778 | significant |
| *Coccidula rufa* | 791 | 1.1005 | significant |
| *Scymnus suturalis* | 1124 | 1.0959 | significant |
| *Amida nigropectoralis* | 810 | 1.1510 | significant |
| *Amida quinquefasiata* | 823 | 1.1351 | significant |

RSF = Relative Simplicity Factor, SSM = Significantly Simple Motifs

### 2.3 Data analysis

The program BLAST (Altschul et al., 1990), National Center for Biotechnology was employed to identify similarities between the isolated sequences and previously published data. The resulting chromatograms of direct sequenced were edited manually by comparing and aligning both strands using SeqMen, a subprogram of DNAstar (DNAstar Inc., 2014). Sequence similarities were assessed using progressive sequence alignment logarithms, alignment was generated using a subprogram of Mega 4.0, the CULSTAL W program (Tamura et al., 2007), using a range of gap open and gap extension penalties. Further analyses were carried out in two major sections; first analyzing Simple repeats, and second analyzing the phylogenetic inferences.

The sequences were analyzed to observe the presence of simple repetitive motifs, following the method of von der Schulenburg et al. (2001), applying two methods to detect the repetitions. The presence of simple repetition and the extent were analyzed by obtaining the Relative Simplicity Factor and Graphical representations, following the methods of Tautz et al. (1986) and von der Schulenburg et al. (2001) with slight modifications, using the computer software SIMPLEv 4.0 (modified version of SIMPLE 34 and SIMPLE 4 of Hancock and Armstrong, 1994). Dot plots were created for each sequence using Gepard 1.19 (Krumsiek et al., 2007) after subjecting the sequences to Dot-matrix analysis to confirm the repetitive regions across the spacer.

The presence and extent of simple repetition were analyzed following the method of Tautz et al. (1986) and von der Schulenburg et al. (2001), using the SIMPLE v 4.0 (modified version of SIMPLE 34 and SIMPLE 4 of (Hancock and Armstrong, 1994) computer software. The program calculates the distribution of tri- and tetranucleotide repeats within a sequence by evaluating the occurrences of each motif within a window of 32 bp 5’ and 3’ to it. Each window is scored for occurrences of the tri- and tetranucleotide motifs at the center by awarding four points for each occurrence of the tetranucleotide motif and three points for each trinucleotide motif (Tautz et al., 1986). Sequence simplicity plots reflect the distribution of these scores along the sequence, with peaks reflecting regions of high concentrations of particular motifs. The program calculates the frequency of motifs according to the following logarithm.

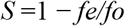

S - Significant threshold
*fe* - average number of times (over 10 sequences) a simplicity score (SS) of this magnitude is observed in the random sequence.
*fo* - observed frequency of the given SS in the test sequence.

Motifs associated with windows whose scores occur at frequencies that exceed the average frequency for that score in the random sequences by a factor of 10 or more are then identified as significantly simple motifs (SSMs), motifs that are clustered at particular positions within the sequence (Hancock and Armstrong, 1994). To compare frequencies of SSMs between species, frequencies of SSMs for a species were normalized to the total number of SSMs in a genome to produce values between zero and one. Using these values mean frequencies of each SSM across species were calculated. Then the correlation coefficients were calculated for each species to compare average frequencies between species. Graphical presentation of SSM for each species was obtained with total ITSI length region and distribution of SSMs across the spacer region observed with the presence of significant simple repeat peaks. The graphical representations of SSMs frequency relevant for each sequence were compared with each dot plot analysis for the presence of repeat motifs.

### 2.4 Phylogenetic analyses

The aligned sequences were analyzed using maximum parsimony (MP), Neighbor-joining (NJ) analyses, and Basiyan likelihood Methods. The resulting chromatograms were edited manually by comparing both strands for taxa using DNAstar – Seqmen subprogram. The computer program (MODELTEST version 3.06; Posada and Crandall, 1998) was used as a subprogram of PAUP*4.0b4a (Swoford, 2002) to select the most adequate substitution model for analyses. The MP was performed using PAUPv 4.0b10 (Swofford, 2002). Heuristic searches were conducted using tree-bisection-reconnection (TBR) branch swapping, 1000 random-addition replicates, and a Max Tree’s value of 1000. Moreover, gaps constitute valuable phylogenetic information (Giribet and Wheeler, 1999) and were treated as a fifth character in the parsimony analyses. Then, the topologies obtained were compared to those found with the gaps treated as missing data. The robustness of nodes was estimated by a bootstrap procedure (Felsenstein, 1985) with 1000 replicates (full heuristic search) of 100 random-addition replicates each, for all analyses. Moreover, in the absence of heterogeneity in the data, adding more data from distinct sources generally increases the accuracy of phylogenetic estimates (Bull et al. 1993; Huelsenbeck et al. 1996), even if several sequences are missing as the benefits of including taxa with missing data in phylogenetic analyses usually overcomes the associated disadvantages (Wiens, 2003; 2005; 2006).

Basiyan likelihood Method was performed with Mr. Bayer’s revolutionary software, starting from a random tree using the program’s default values for the prior probabilities, consisting of four simultaneous Markov chains, three heated and one cold, which were run for 10^6^ generations. The Bayesian likelihood analyses were repeated four times and a single tree was sampled randomly every 100th generation. The log-likelihood scores for all generations were studied to identify the burn-in phase of initial generations in which likelihood scores progressively improve until they fluctuate narrowly around a stable value. In all Bayesian analyses, the latter 5000 sampled trees from each analysis were pooled (after confirming they had converged on similar log-likelihood values) and imported into PAUP* 4.0b4a and a 50% consensus tree computed, with the support values for each branch constituting their posterior probability. Clades with posterior probabilities >95% were considered as significantly supported by the data (Huelsenbeck et al., 2002).

Further, the evolutionary history was inferred using the Neighbor-Joining (NJ) method (Saitou and Nei, 1987). The bootstrap consensus tree was from 1000 replicates (Felsenstein, 1985) is taken to represent the evolutionary history of the taxa analyzed (Felsenstein, 1985). Branches reproduced in less than 50% bootstrap replicates were collapsed. The percentage of replicate trees are shown adjacent to the branches in which the associated taxa clustered together in the bootstrap test (1000 replicates) (Felsenstein, 1985). In pairwise sequence comparisons, all positions containing alignment gaps and missing data were eliminated. Phylogenetic trees view with the program TREEVIEW (win 32), version 1.6.6 (Page, 2001).

## 3.0 Results

### 3.1 Length Variation of Internal Transcribed Spacer I

The amplified ITSI region, including the 5.8S and 18s ribosomal subunits, are between 791–1671 bases in length, which is 791 bp minimum in *Coccidula rufa* and maximum is in *Exochomus quadripustulatus,* having 2572 bases. The primers amplified two adjacent regions; 18S with a length of 44 bp and 5.8 with a length of 112 bp when compared with the published sequences in Genbank. All sequenced ITS 1 gene region was larger than the previously studied ITSI of other polyphagous beetles, showing lengths >550 bp. The size of the ITSI region drastically varies inter specifically, including closely related taxa of the same subfamily or tribe. However, observed intra-specific sequence length variability was only with a few bases. In the present study ITSI gene regions sequenced from *Psylloborus vigintictopunctata, Coccinella spetempunctata, Harmonia axyrid,is* and *Adalia bipunctata* were compared with the existing sequences deposited in the NCBI. All the comparisons indicated the absence of intraspecific variability. The differences observed were; *Adalia bipunctata* showed a 3 bp maximum size difference and *Psyllobora vigintiduopunctata* showed a 1 bp size difference (Table 02).

The base composition of the ITSI region of Coccinellidae showed highly similar values of A, T, G and C (Fig 01). *Amida nigropectoralis* and *Amida quingquefasiata* has the lowest A+T percentage of 47.1 where total base composition is 24.6% T, 25.8% C, 22.5% A, 27.1% G for *Amida quingquefasiata* and 24.3% T, 26.4% C, 22.8% A, 26.5% G for *Amida nigropectoralis. Phyllobora vigintiduopunctata* showed the highest A+T percentage of 55.6 where the total base composition is 25.6% T, 24.3% C, 30.0% A, 20.1% G. The outgroup *Diabrotica barberi* and *Cerotoma trifurcata* of family Chrysomelidae showed a comparatively higher percentage of A+T, 63.7 and 64.1% respectively with the total base composition values of 29.6% T, 19.8% C, 34.1% A, 16.4% G in *Diabrotica barberi* and 31.9% T, 18.7% C, 32.2% A, 17.2% G in *Cerotoma trifurcate. Diabrotica barberi* and *Cerotoma trifurcata* showed similarity to analyze sequences only at the region of 5.8S of ribosomal DNA having 56 bases.

**Fig. 01:**
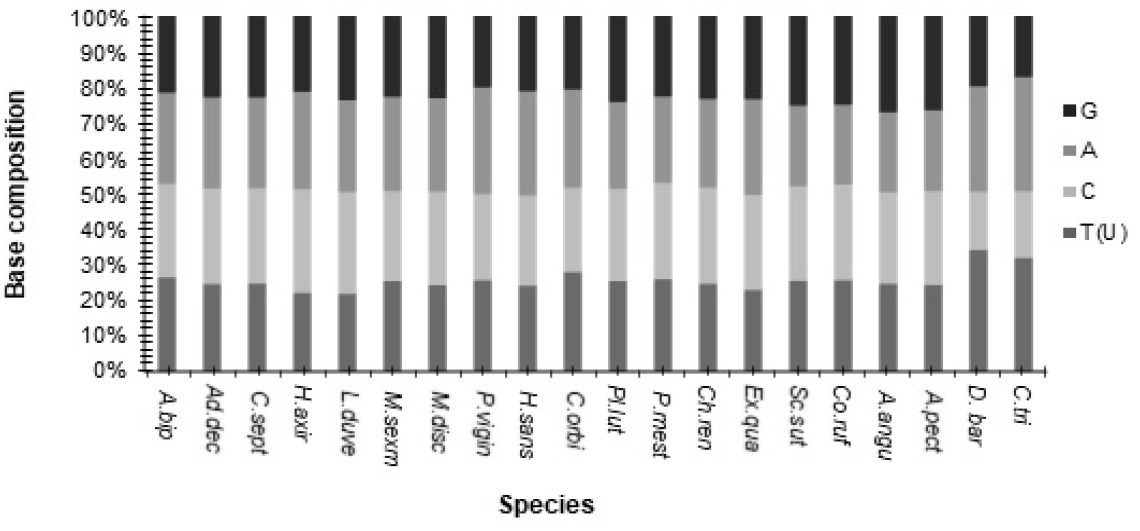
The base composition of ITSI regions; *bip* - *Adalia bipunctata, A. dec - Adalia decempunctata, C. sep - Coccinella septempunctata, H. axy - Harmonia axyridis, L. duv - Lemnia duvauceli, M. sex - Menochilus sexmaculata, M.dis - Micraspis discolor, P. vig - Phyllobora vigintiduopunctata, H. san - Halyzia sanscrita, P. lut - Platynaspis luterubra, P. mes - Phaenochilus mesternalis, C. ren - Chilocorus renipustulatus, E. qua - Exochomus quadripustulatus, S. sut - Scymnus suturalis, C. ruf - Coccidula rufa, A. qui - Amida quingquefasiata, A. nig - Amida nigropectoralis, D. ba - Diabrotica barberi, C. tri – Cerotoma trifurcate*

### 3.2 Presence of simple repetition

The detailed analyzed results of the SIMPLE v 4.0 programs are summarized in table 04 with the Relative Simplicity Factor – RSF and the presence of Significantly Simple Motifs – SSMs for each species. Except for *Harmonia axyridis, Phyllobora vigintiduopunctata, Chilocorus renipustulatu, Lemnia duvauceli* all other species showed a significant number of simple repetitions (Table 04). Among them, *Adalia bipunctata* showed the highest simple repetition with the RSF value of 1.4440 which was similar to the von der Schulenburg *et al.* (2001) records and the lowest significant RSF value is 1.0891 for *Halyzia sanscrita.*

**Table 04:** Frequencies of significant simple motifs (SSMs) of different species.

|  | ABIP<sup>a</sup> | ADEC | CSEP | MDIS | MSEX | HSAN | EQUA | PMES | PLUT | SSUT | AANG | APEC | CRUF | Average |
| --- | --- | --- | --- | --- | --- | --- | --- | --- | --- | --- | --- | --- | --- | --- |
| CGG | 0.015 | 0.000 | 0.000 | 0.000 | **0.889** | 0.000 | 0.000 | 0.000 | 0.000 | 0.000 | **0.429** | **0.438** | **0.167** | **0.108** |
| CGC | **0.052** | **0.227** | 0.000 | 0.000 | 0.000 | 0.000 | 0.000 | 0.000 | **0.250** | **0.111** | **0.214** | **0.188** | **0.167** | **0.067** |
| CGT | **0.125** | **0.205** | **0.400** | **0.333** | 0.000 | 0.000 | **0.224** | **0.250** | **0.250** | **0.056** | **0.214** | **0.250** | **0.500** | **0.163** |
| AAC | 0.000 | 0.000 | 0.000 | 0.000 | 0.000 | 0.000 | 0.000 | 0.000 | 0.000 | 0.000 | **0.143** | **0.063** | 0.000 | **0.067** |
| ATA | **0.140** | 0.000 | 0.000 | 0.000 | 0.000 | 0.000 | **0.224** | 0.000 | 0.000 | 0.000 | 0.000 | 0.000 | 0.000 | 0.016 |
| AAT | **0.140** | 0.000 | 0.000 | 0.000 | 0.000 | 0.000 | 0.000 | 0.000 | 0.000 | 0.000 | 0.000 | **0.063** | 0.000 | 0.011 |
| TCG | **0.096** | **0.227** | **0.400** | 0.000 | 0.000 | **0.300** | **0.207** | 0.000 | 0.000 | **0.389** | 0.000 | 0.000 | **0.167** | 0.020 |
| TAA | **0.090** | 0.000 | 0.000 | 0.000 | 0.000 | 0.000 | **0.052** | 0.000 | 0.000 | 0.000 | 0.000 | 0.000 | 0.000 | 0.011 |
| CGA | **0.081** | **0.227** | **0.100** | 0.000 | 0.000 | **0.600** | 0.017 | 0.000 | 0.000 | **0.333** | 0.000 | 0.000 | 0.000 | **0.103** |
| ATG | **0.052** | 0.000 | 0.000 | 0.000 | 0.000 | 0.000 | 0.035 | 0.000 | 0.000 | 0.000 | 0.000 | 0.000 | 0.000 | 0.007 |
| GAG | **0.052** | 0.000 | 0.000 | **0.167** | 0.000 | 0.000 | 0.000 | 0.000 | 0.000 | 0.000 | 0.000 | 0.000 | 0.000 | **0.094** |
| GAA | 0.044 | 0.000 | 0.000 | 0.000 | 0.000 | 0.000 | 0.000 | **0.250** | **0.083** | 0.000 | 0.000 | 0.000 | 0.000 | 0.004 |
| AAA | 0.022 | **0.091** | 0.000 | 0.000 | 0.000 | 0.000 | 0.000 | 0.000 | **0.167** | 0.000 | 0.000 | 0.000 | 0.000 | 0.016 |
| GAT | 0.015 | 0.000 | 0.000 | **0.167** | 0.000 | **0.100** | 0.000 | 0.000 | 0.000 | 0.000 | 0.000 | 0.000 | 0.000 | 0.026 |
| TAT | 0.007 | 0.023 | 0.000 | 0.000 | 0.000 | 0.000 | **0.138** | 0.000 | 0.000 | 0.000 | 0.000 | 0.000 | 0.000 | 0.001 |

**Table 04: Frequencies of significant simple motifs (SSMs) of different species (Continued...)**
|  | ABIP<sup>a</sup> | ADEC | CSEP | MDIS | MSEX | HSAN | EQUA | PMES | PLUT | SSUT | AANG | APEC | CRUF | Average |
| --- | --- | --- | --- | --- | --- | --- | --- | --- | --- | --- | --- | --- | --- | --- |
| AAG | 0.007 | 0.000 | 0.000 | 0.000 | 0.000 | 0.000 | 0.000 | 0.000 | 0.000 | 0.000 | 0.000 | 0.000 | 0.000 | 0.040 |
| GAC | 0.007 | 0.000 | **0.100** | 0.000 | 0.000 | 0.000 | 0.000 | **0.500** | 0.000 | 0.000 | 0.000 | 0.000 | 0.000 | 0.001 |
| ATC | 0.007 | 0.000 | 0.000 | 0.000 | 0.000 | 0.000 | 0.000 | 0.000 | 0.000 | 0.000 | 0.000 | 0.000 | 0.000 | 0.001 |
| TAC | 0.000 | 0.000 | 0.000 | 0.000 | **0.111** | 0.000 | 0.000 | 0.000 | 0.000 | 0.000 | 0.000 | 0.000 | 0.000 | 0.006 |
| GTA | 0.007 | 0.000 | 0.000 | 0.000 | 0.000 | 0.000 | 0.000 | 0.000 | 0.000 | 0.000 | 0.000 | 0.000 | 0.000 | 0.001 |
| AAT | 0.044 | 0.000 | 0.000 | 0.000 | 0.000 | 0.000 | 0.000 | 0.000 | 0.000 | 0.000 | 0.000 | 0.000 | 0.000 | 0.002 |
| GTT | 0.000 | 0.000 | 0.000 | **0.333** | 0.000 | 0.000 | 0.000 | 0.000 | 0.000 | 0.000 | 0.000 | 0.000 | 0.000 | 0.020 |
| GTC | 0.000 | 0.000 | 0.000 | 0.000 | 0.000 | 0.000 | 0.000 | 0.000 | **0.250** | **0.111** | 0.000 | 0.000 | 0.000 | 0.022 |
| GTG | 0.000 | 0.000 | 0.000 | 0.000 | 0.000 | 0.000 | 0.017 | 0.000 | 0.000 | 0.000 | 0.000 | 0.000 | 0.000 | 0.001 |
| ACG | 0.000 | 0.000 | 0.000 | 0.000 | 0.000 | 0.000 | 0.000 | 0.000 | 0.000 | 0.000 | 0.000 | 0.000 | 0.000 | 0.002 |
| ATT | 0.000 | 0.000 | 0.000 | 0.000 | 0.000 | 0.000 | 0.035 | 0.000 | 0.000 | 0.000 | 0.000 | 0.000 | 0.000 | 0.001 |

<sup>a</sup>SSM frequencies for each group are represented as a proportion of the total. Values greater than 0.05 are in bold. Zero values are in italics.
ABIP - *Adalia bipunctata*, ADEC - *Adalia decempunctata*, CSEP - *Coccinella septempunctata*, MDIS - *Micraspis discolor*, MSEX - *Menochilus* *sexmaculata*, HSAN - *Halysia sanscrita*, EQUA - *Exochomus quadripustulatus*, PMES - *Phaenochilus mesternalis*, PLUT - *Platynaspis luterubra*, SSUT - *Scymnus suturalis*, AANG - *Amida quinguefasiata*, APEC - *Amida nigropectoralis*, CRUF - *Coccidula rufa*

The dot plot analyzes of each sequence were studied with the graphical SSM presentation as a comparison. The regions with high repeatability are shown in the dot matrix with a high stringent area. *Harmonia axyridis* generally have lower levels of cryptically significant regions with no detectable SSMs. The dot plot and graphical SSMs comparative analyses represent equally low dense regions comparative to other detectable SSM species. All other species with higher significant SSM regions; *Adalia bipunctata, Adalia decempunctata, Menochilus sexmaculata, Coccinella septempunctata, Micraspis discolor, Halyzia sanscrita, Phaenochilus mesternalis*, *Platynaspis luterubra*, *Exochomus quadripustulatus*, *Coccidula rufa, Scymnus suturalis, Amida nigropectoralis and Amida quingquefasiata* represent the same analyses in dot matrix presentation also, with the presence of high stringent areas.

Table 04 explains the frequency of different SSMs of different species as measured by the program. The most common motifs calculated as the average of all species were CGG, CGC, CGT, AAC, CGA, and GAG all of which average more than 0.05 of all SSMs. *Adalia bipunctata* is having the most SSMs and the highest density of SSMs, and the second *Adalia decempunctata.* Compared to the SSM content averaged for overall species, SSM contents of individual species showed a range of correlation coefficients (Table 05). The strongest (*r* > 0.861) were seen for *Coccidula rufa. Phaenochilus mesternalis* shows a narrower correlation coefficient (r > 0.300).

**Table 05:** Correlation coefficients between species SSMs frequencies and corresponding domain average.

| Species | $r^a$ |
| --- | --- |
| *Adalia bipunctata* | 0.444 |
| *Adalia decempunctata* | 0.760 |
| *Menochilus sexmaculata* | 0.362 |
| *Coccinella septempunctata* | 0.783 |
| *Harmonia axyridis* | N/A<sup>b</sup> |
| *Lemnia duvauceli* | N/A |
| *Micraspis discolor* | N/A |
| *Phyllobora vigintiduopunctata* | N/A |
| *Halyzia sanscrita* | 0.440 |
| *Chilocorus renipustulatus* | N/A |
| *Phaenochilus mesternalis* | N/A |
| *Platynaspis luterubra* | 0.418 |
| *Exochomus quadripustulatus* | 0.494 |
| *Coccidula rufa* | 0.861 |
| *Scymnus suturalis* | 0.575 |
| *Amida nigropectoralis* | 0.675 |
| *Amida quinguefasiata* | 0.631 |

<sup>a</sup>Correlation coefficient between the SSM frequencies seen in the given species and the mean frequencies given in Table 04
<sup>b</sup>N/A=Species had no SSMs

The significant repeat motifs could be identified with high peaks and motif/s. As this method reveals statistically significant regions of simple sequence DNA as it occurs either in tandemly repeated arrays; pure simple sequence DNA, (Tautz and Dover, 1986), or in regions in which a few repetitive motifs are scrambled (Cryptic simplicity). The graphics displays of high mountain peaks indicate that significant species having almost all regions are cryptically simple through the repetition of one short motif or another. The motifs associated with these peaks or the SSM (Hancock and Armstrong, 1994) are indicated some regions can contain a number of SSMs: such regions probably contain tandem arrays of a single SSM. The graphical representations showed interesting results about the distribution of SSMs among species. *Adalia bipunctata, Adalia decempunctata, Exochomus quadripustulatus* shows high accumulation of AT content repeats in 5’ flanking region, while high accumulation of GC repeats in 3’ flanking region. *Amida nigropectoralis, Amida quingquefasiata, Menochilus sexmaculata, Phaenochilus mesternalis, Coccidula rufa, Platynaspis luterubra s*hows higher concentration of GC repeats towards the 3’ flanking region. In the sequences where significant number of simple repeats are present, the middle of the spacer is free of SSM or has very few compared to the flanking regions.

### 3.4 Phylogenetic analyses of Internal Transcribed Spacer Region I

The analyses of individual Internal Transcribed Spacer I regions with three different methods yielded the almost same tree topology (Fig. 02). The same tree topology was obtained by von der Schulenburg *et al.* (2001) with species from four subfamilies of the Coccinellidae. The tribe Psylloborini of subfamily Coccinellidae was monophyletic. The members of the Subfamily Chilocorinae first associate with the tribes of subfamily Coccinellidae and then relate with the tribe Scymnini. *Exochomus quadripustulatus* which shows the highest length variably showed a relationship with *Phaenochilus mesternalis* 77 bootstrap and 84 posterior probabilities. *Platynaspis luterubra* showed 100 bootstrap relationships with *Scymnus suturalis.* The close relationship of tribe Coccidulini with Ortaliini is prominent in the present analyses.

**Fig. 02:**
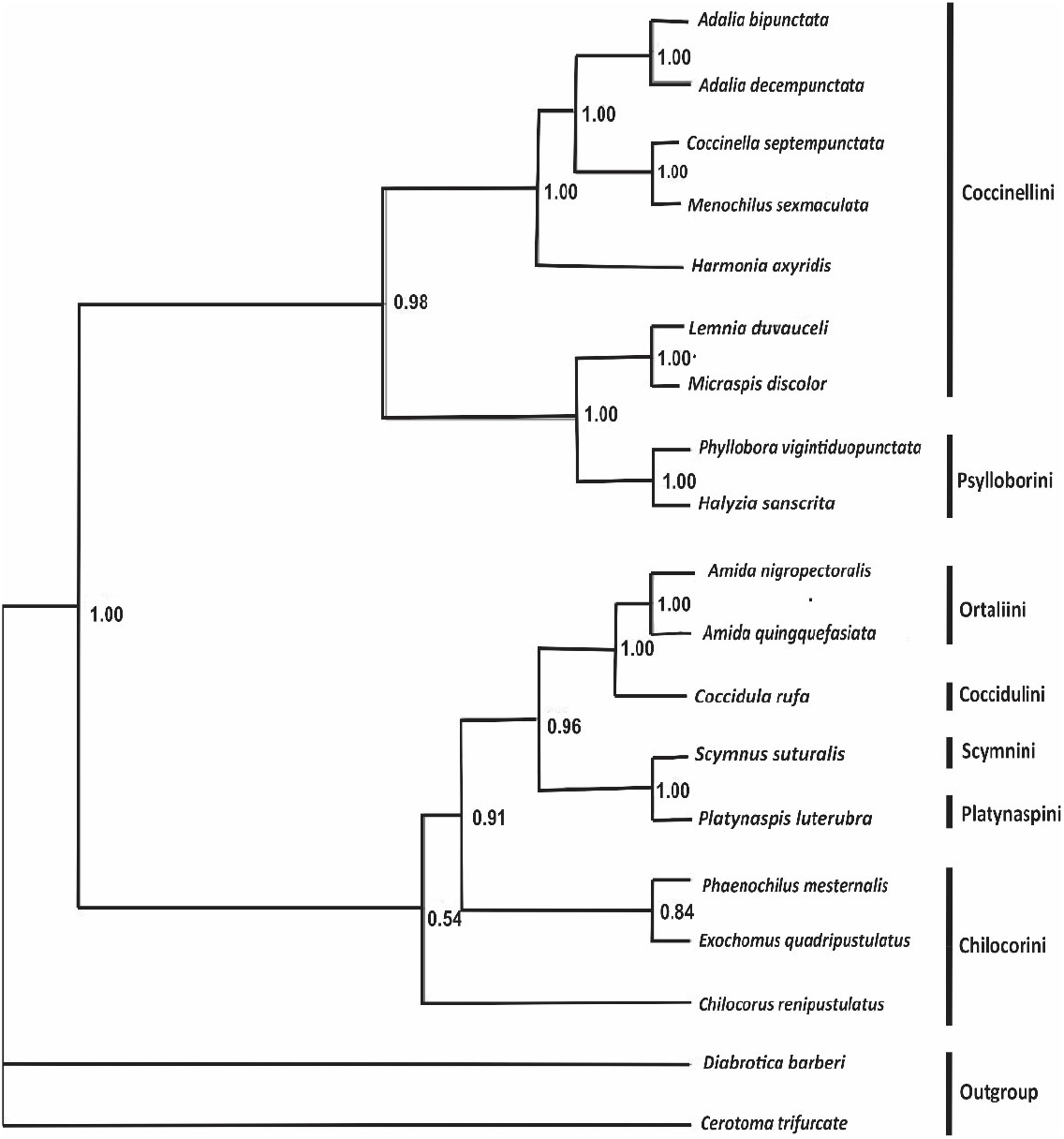
Mr Bayes Posterior bootstrap phylogeny inferred from ITSI gene region. Posterior probability values are shown near the branches.

## 4.5 Discussion

The isolated Internal Transcribed Spacer regions of Coccinellidae species are comparatively longer than the previously studied ITSI regions, length differences could be observed among genera, species, tribes as well as species levels. These results are similar to the previous study done by von der Schulenburg *et al.* (2001), who first reported the extreme length and length variation of ITSI region of Coccinellidae species. He recorded the unique nature of sequence variability of ITSI spacer region among species, genera, tribes, and subfamilies. The majority of Eukaryotes usually show ITSI lengths of no more than 800 bp, and almost all of them are shorter than 1,100 bp (Schlötterer et al., 1994; Vogler and DeSalle, 1994; Bowles et al., 1995; McLain et al., 1995; Miller et al., 1996; Tang et al., 1996; Fenton et al., 1997; Downie et al., 1998; Gouliamova and Hennebert, 1998; Kumar, et al., 1999; van Herwerden, et al., 1999; von der Schulenburg et al., 2001; Harris and Crandall, 2000). Very recently Seu Ki et al. (2007) reported extremely long PCR fragments of ITSI and 5.8S genes from *Cochlodinium polykarikoides* compared to other Dinoflagellates. In addition to this report there are only two documented exceptions written in the history of ITSI length variation; *Schistosoma japonica* (Platyhelminthes) and the *Anopheles gambiae* (Insecta) species complexes, which produce maximum ITS1 sizes of 1,400 and at least 5,500 bp, respectively (Paskewitz et al., 1993; van Herwerden et al., 1998). In addition, size variation between closely related taxa is usually less than 400 bp, except for the three above-mentioned cases, which yield length differences of about 900 and at least 3,000 bp.

von der Schulenburg *et al.* (2001) studied the sequences extensively to find the similar regions and long repeat units which are responsible for extreme length variation of ITSI spacer. He reported long tandemly repeated sequence motifs of greater than 60 bp in *Adalia bipunctata, Adalia decempunctata, Harmonia axyridis, Phyllobora vigintiduopunctata Chilocorus renipustulatu, Exochomus quadripustulatus and Coccinella septempunctata.* The presence of long repeats is confined to the middle of the spacer of all the species. According to von der Schulenburg et al. (2001) the extreme size and size variation seems to be associated with long repetitive elements. However, they reported simple repeats only in *Adalia bipunctata.* The present study resulted in eleven species with a significant number of simple repeat motifs out of seventeen species studied. Interestingly *Harmonia axyridis,* P*hyllobora vigintiduopunctata, Coccinella septempunctata, Chilocorus renipustulatus,* which were reported with long repetitive sequences by von der Schulenburg et al. (2001) did not show any significant simple repeats (SSM). However, *Exochomus quadripustulatus, Adalia bipunctata, Adalia decempunctata* the species having long internal repeats, were present with simple repeats also.

The dot matrix analysis, and their attendant SSMs need to be considered together to interpret the nature of the repetitive structure of a given sequence. The dot plot analyses and the SSMs graphical representation give a clear idea about the presence of simple repeats by tallying the high stringent areas and graphical peaks of the repetitive regions. Sequence analysis of the data from coccinellids indicated an important factor, which characterizes ITS1 evolution in this group. In particular, size variation between coccinellid species was associated with a high degree of sequence divergence. For all taxa and almost all subsets of the data, simple repeats represent a specific distribution pattern by accumulating higher level peaks of SSM in graphs and by observing high stringent areas in dot plots. These high abundances repeat regions were confined to either of the flanking regions except other than in *Micraspis discolor.* Almost all of them were having repetitive regions in 3’ flanking region. Hence, sequence simple repetitions are present only in restricted areas, suggesting that they are due to functional constraints. This seems likely for the small regions towards the flanking regions of the spacer, such regions have been reported previously associate with secondary structure motifs having functional importance. Earlier studies indicated that conserved secondary-structure motifs in ITSI region were primarily found at the 3’ and 5’ ends in yeast to contain various elements required for ribosomal biogenesis (van Nues et al., 1994; Weaver et al., 1997). For metazoans functional importance has been implicated at least for 3’ end (Schlötterer et al., 1994; von der Schulenburg et al., 2001).

In addition, our analysis indicates that significantly simple repetitive regions are mainly confined with a high amount of G/C content when refer the mean frequencies of motifs. Recently Seu Ki et al., 2007 studied on *Cochlodinium polykrikoides* and reported ITSI having considerably high G/C content, particularly in repeat units that are considered structurally stable, playing an important role in ribosome biogenesis through the processing of the pre-RNA. Moreover, in the base composition analysis of the ITSI regions of Coccinellidae species, all four bases were found in equal frequencies. This suggests indeed, that the repeat regions are having an accumulation of G/C bases across the spacers. These observations, taken together, suggest that simple repetitions in the Coccinellinae ITSI are representing functional regions basically in the 3’ flanking region. In addition, the present study included data for more than two species from the same genus; eg. *Adalia* spp, and *Amida* spp. in comparison of graphical SSMs regions across genus level apparent repeat similarities were observed. However, the frequency of occurring motif repeatability shows some difference between two species of both genera (Table 05). Moreover, for two subfamilies, species were included from more than one tribe; Chilocorinae and Coccinellinae. *Phyllobora vigintiduopunctata* does not show any significant simple repetitions in tribe Psylloborini of subfamily Coccinellinae, whereas *Halyzia sanscrita* presence with significant simple repetitions. This is true for the Chilocorinae, with the presence of significant number of simple repeats in *Exochomus quadripustulatus* from the studied two tribes. Taken together all the above observations suggest that sequence simple repetition in Coccinellidae does not share show similarities above the genera level, however, the similarity of simple repetitive regions is closer to the species-specific or sometimes specific among a closely related group of species only.

In addition, AT-rich simple repeats are also present in *Adalia bipunctata, Adalia decempunctata and Exochomus quadripustulatus*. Generally, AT-rich regions are more unstable than GC-rich regions. If several regions in a gene (or even unlike parts of the genome) are affected by these mutational processes, a bias in AT content, degree of repetition, and sequence composition should arise. This possibility would also explain a common AT bias in different length-variable regions of rRNA molecule, as an example, AT-rich slippage-generated motifs have occurred during the evolution of *Drosophila melanogaster* rDNA (Hancock and Dover, 1988). The phylogenetic assessment of the ITSI gene region was successful to obtain a robust support tree. Sequence and length variation as well as the nuclear ribosomal repeats of the ITS I region provides information on phylogenetic evidence among closely related taxa, species identification, and population studies. The ITS region has generally been used in phylogenetic studies of plants and fungi (Lohtander et al., 2000; Lantz et al., 2002), although until recently, comparatively few insect studies have focused on this genetic marker (Schlötterer et al., 1994, von der Schulenburg *et al.*, 2001, Gallego *et al.*, 2001, Seu Ki *et al.*, 2007). An independent examination of this important gene permits the tree topology from all three methods studied and the tree topology was similar to the previous phylogenetic analyses. The position of Coccinellidae in the tree is clear. The tribe Phylloborini represents its position clearly within the subfamily Coccinellidae. Subfamily Chilocorinae is closely linked with Coccinellinae and then to the Subfamily Aspidemerinae, Coccidulinea and Ortalinea show phylogenetically older toxon levels showing similar results.

